# Single-component optogenetic tools for inducible RhoA GTPase signaling

**DOI:** 10.1101/2021.02.01.429147

**Authors:** Erin E. Berlew, Ivan A. Kuznetsov, Keisuke Yamada, Lukasz J. Bugaj, Joel D. Boerckel, Brian Y. Chow

## Abstract

We created optogenetic tools to control RhoA GTPase, a central regulator of actin organization and actomyosin contractility. RhoA GTPase, or its upstream activating GEF effectors, were fused to BcLOV4, a photoreceptor that can be dynamically recruited to the plasma membrane by a light-regulated protein-lipid electrostatic interaction with the inner leaflet. Direct membrane recruitment of these effectors induced potent contractile signaling sufficient to separate adherens junctions in response to as little as one pulse of blue light. Cytoskeletal morphology changes were dependent on the alignment of the spatially patterned stimulation with the underlying cell polarization. RhoA-mediated cytoskeletal activation induced YAP nuclear localization within minutes and subsequent mechanotransduction, verified by YAP-TEAD transcriptional activity. These single-component tools, which do not require protein binding partners, offer spatiotemporally precise control over RhoA signaling that will advance the study of its diverse regulatory roles in cell migration, morphogenesis, and cell cycle maintenance.

## INTRODUCTION

RhoA, a member of the Rho-family of small GTPases, centrally regulates actin organization and actomyosin contractility in cell migration, cell cycle maintenance, and developmental morphogenesis^1,2^. Key amongst its diverse roles, RhoA signaling dynamics coordinate actin stress fiber formation that determines how cells generate cytoskeletal tension to transmit mechanical forces across the cell, across neighboring cell-cell junctions, and to the extracellular matrix (ECM) via focal adhesions^3^. Thus, new tools for inducible control over RhoA activity may greatly enhance understanding of cytoskeletal dynamics and mechanotransduction, the cyto-mechanical activation of gene transcription^4,5^.

Optogenetics is highly attractive for these purposes owing to its high spatiotemporal precision vs. pharmacological and genetic techniques, which can be encumbered by slow uptake/washout kinetics and frequent pleotropic effects. Because small GTPases and their activating GEFs (guanine nucleotide exchange factors) signal at the plasma membrane, optogenetic membrane localization techniques are effective for inducible control over their function, where cytosol-sequestered effector proteins are dynamically recruited to the cytosol-facing inner leaflet of the plasma membrane to upregulate effector signaling^6^. Based on earlier reported chemically inducible dimerization (CID)-based approaches for RhoA membrane recruitment^7,8^, heterodimerization between a photosensory protein and a protein binding partner (one of which is membrane localized) has been widely used to control upstream RhoA-activating GEFs^9–13^ and phosphatases^14^. This strategy is sensitive to the expression level and stoichiometry of the two components, and thus may require expression level-tuning by visualizing multiple fluorescent reporters and/or clonal cell line selection^15–17^. Previously, we reported the direct optogenetic control over RhoA GTPase itself, where inducible cryptochrome clustering presumably increases the binding avidity of the GTPase to GEFs. However, this system has limited spatial resolution due to cytosolic diffusion beyond the optical stimulation field as it oligomerizes prior to stable membrane localization^18^.

Recently, we reported that BcLOV4, a light-oxygen-voltage (LOV) flavoprotein from *Botrytis cinerea*, rapidly translocates to the plasma membrane in mammalian cells via a blue light-regulated electrostatic protein-lipid interaction (PLI) with the inner leaflet^20,21^. This direct interaction with the membrane itself is powerful for creating “single-component” tools for dynamic membrane recruitment of peripheral membrane proteins, without the obligate heterodimerization- or self-oligomerization protein partners of the aforementioned (PPI) proteinprotein interaction-based systems. We previously leveraged the intrinsic membrane-binding capability of BcLOV4 fused to Rho-family Cdc42-GEF and Rac1 GTPase proteins to induce filopodial and lamellipodial protrusions^6,22^. Here, we report the engineering of single-component optogenetic RhoA GTPase and GEFs to potently drive actomyosin contractility, stress fiber formation, and rapid activation of transcriptional mechanotransduction (**Figure 1a, Supplementary Figure 1**).

**FIGURE 1:**
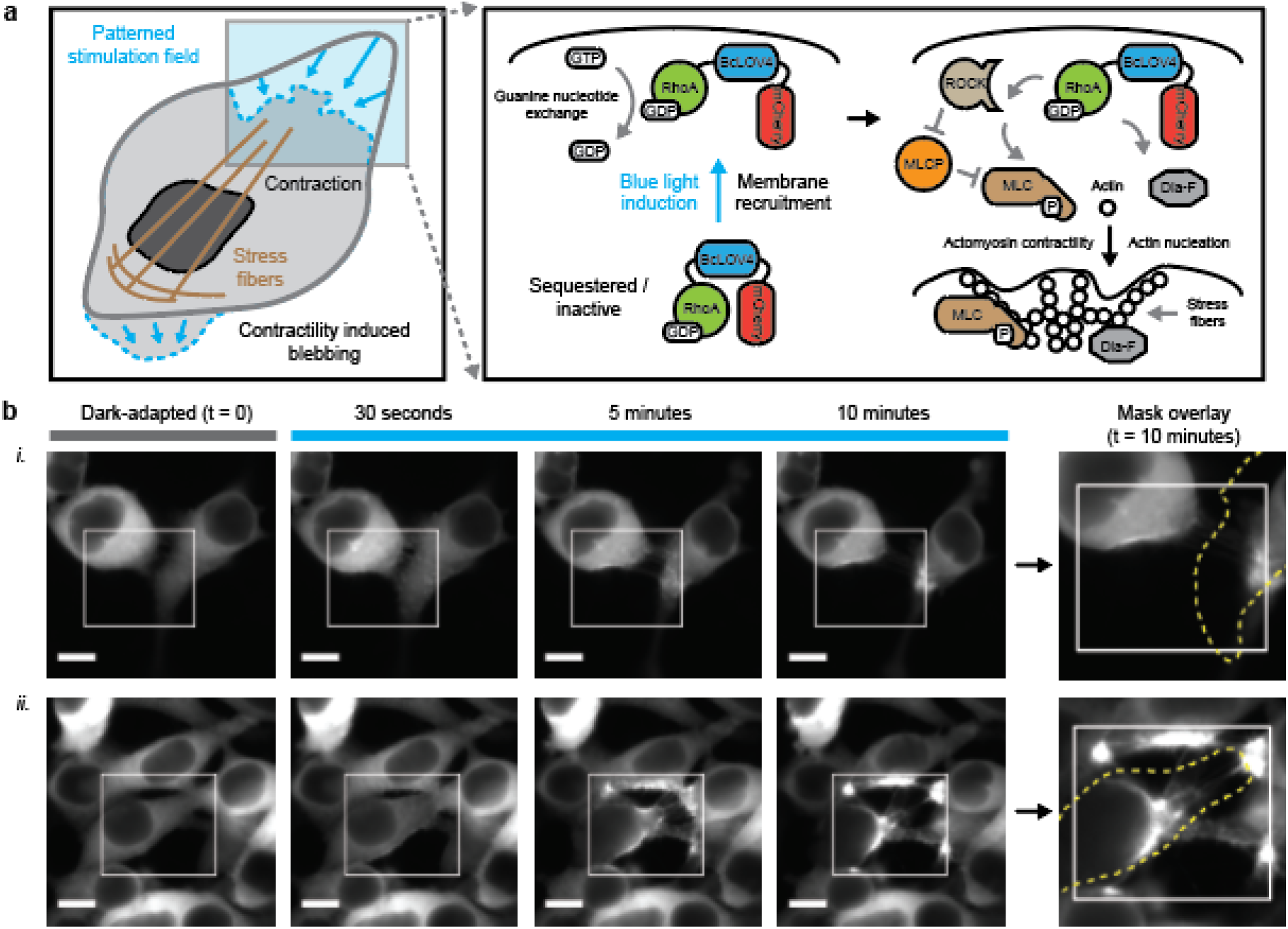
Single-component optogenetic control over RhoA signaling. **a**. Schematized induction of cytoskeletal changes and contractile signaling in response to opto-RhoA activation by dynamic membrane recruitment mediated by the light-regulated electrostatic protein-lipid interaction of BcLOV4 with the plasma membrane. **b**. Epifluorescence micrographs of HEK cells expressing opto-RhoA, visualized by mCherry. i. Trailing edge contact in two adjacent cells. ii. 4-cell adherens junction separation. White box = spatially patterned blue light iillumination field, stimulated at 1.6% duty ratio. Dotted yellow line = cell boundary mask in the dark-adapted state. Scale = 10 μm.

**Supplementary Figure 1: Single-component optogenetic control over RhoA signaling by GEF activation.**

## RESULTS

### Engineering of single-component tools

The genetic constructs were designed and engineered as previously described^22^. We screened domain arrangement combinations of BcLOV4, mCherry fluorescent reporter, and either RhoA or the catalytic DH (Dbl-homology) domain of RhoA-activating ARHGEF11. The domains were each separated by flexible glycineserine-rich (GGGGS)_2_ linkers, and the six respective domain arrangements were tested for their respective expression levels, subcellular distribution characteristics, and inducible translocation efficiency in HEK293T cells (**Supplementary Figures 2-3**), a standard cell line for membrane protein engineering. For opto-RhoA, the most favorable domain arrangement was an N-terminal fusion to BcLOV4-mCherry (RhoA-BcLOV4-mCherry), and in opto-RhoGEF11, it was BcLOV4-RhoGEF11-mCherry.

**Supplementary Figure 2: Molecular engineering of opto-RhoA.**

**Supplementary Figure 3: Molecular engineering of opto-RhoGEF.**

The cytoskeletal response to digital micromirror device (DMD)-patterned illumination was pronounced. Sparse pulsatile stimulation (duty ratio ϕ = 1.6% or 1s per minute; λ = 450 nm at 15 mW/cm^2^) potently induced contractility sufficient to separate intercellular adherens junctions and create membrane blebs from transient delamination of the membrane from the actyomysin network (**Figure 1b**). The duty ratio was initially chosen to ensure that the optogenetic activity was not photochemically limited, by providing one flavin photochemistry-saturating pulse per membrane association-dissociation cycle.

To our initial surprise, a single 5s-pulse of whole-field or unpatterned stimulation was also sufficient to induce extensive morphological changes (**Supplementary Figure 4**). This feature of opto-RhoA and -RhoGEF11 differs from our previous work with single-component Rac1 GTPase or Intersectin Cdc42-GEF, which do not produce appreciable protrusions without DMD patterning or (ROI) region-of-interest selection. The ability to use unpatterned stimulation to drive morphological changes over a whole field-of-view (FOV) facilitated experimental throughput, statistical powering, and blinding for automated data analysis and systematic characterization (**Figure 2a and Supplementary Figure 5**).

**FIGURE 2:**
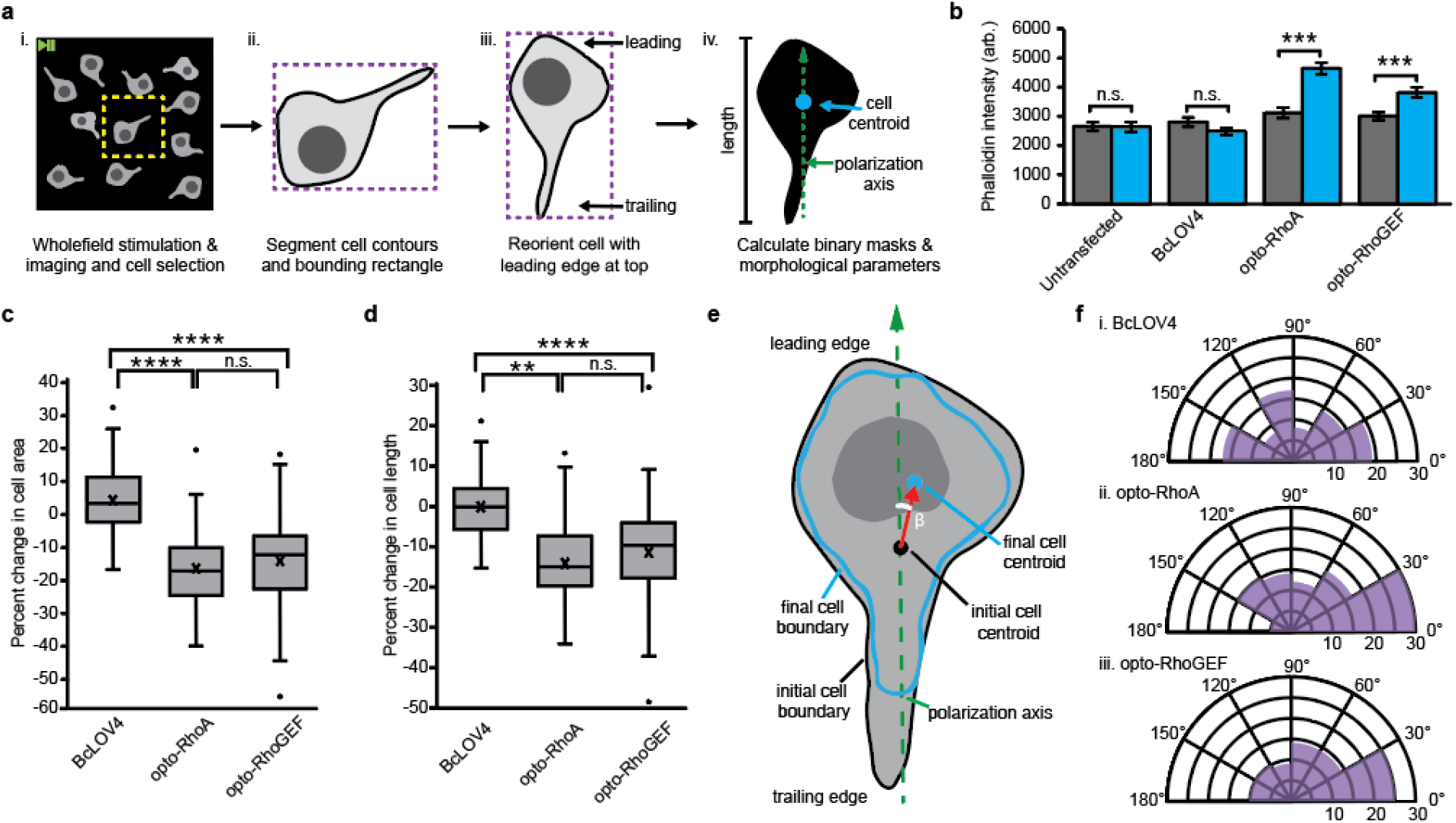
Optogenetic induction of contractility in response to unpatterned wholefield stimulation. **a**. Image analysis workflow. i. Whole-field-of-view timecourse videos are cropped (yellow) to contain only one cell. ii. For the initial frame, cell contours (black) and the bounding rectangle (purple) are identified and defined in OpenCV (threshold function and imutils). iii. The cell is iteratively rotated in 5° increments. The angle that maximizes rectangle height and positions the nucleus closer to the top is applied to all frames so that the y-axis aligns with the cell polarity. iv. Binary masks are created of the cell at initial and final timepoints, from which cell areas, centroids, and lengths are calculated. **b**. Phalloidin stain intensity in dark-adapted vs. stimulated cells. Mean +/- standard error. N = 40 cells per condition. **c**. Box-and-whisker plot of cell area change upon wholefield stimulation. **d**. Box-and-whisker plot of cell length change upon wholefield stimulation. N = 82-93 cells per condition. **e**. Schematized calculation of angle of cell movement. Centroids of the initial (red) and final (blue) cell boundaries are calculated in OpenCV (moments function). The angle of movement between the cell polarization vector (green, dashed) and the centroid movement vector (red) is designated as β. **f**. Circumplex charts of the angle of movement relative to the polarization vector in cells expressing i. BcLOV4, ii. opto-RhoA, and iii. opto-RhoGEF. N = 82-93 cells per condition. **b-d** Mann-Whitney U test: (**) p < 0.01; (***) p < 0.001; (****) p < 0.0001; (n.s.) not significant.

**Supplementary Figure 4: Optogenetic induction of contractility by a single pulse of unpatterned stimulation.**

**Supplementary Figure 5: Actin imaging of HEK cell polarization axis.**

### Systematic characterization of optogenetic activity

As primary measures of morphological dynamics, we quantified the changes in cell area, cell length along the polarization axis, and cell centroid displacement vector upon blue light induction. Stress fiber levels were quantified by phalloidin staining (**Figure 2b**). Optogenetic activation drove cytoskeletal changes that were highly significant and of large effect size vs. the BcLOV4-mCherry control (**Figure 2c-d**). The cell centroid displacement vector was highly preferential toward the leading edge (**Figure 2e-f**), meaning that the cytoskeletal retraction was predominately at the trailing edge and along the polarization axis upon stimulating the whole cell. This trend suggests that the tensile asymmetry introduced by the underlying cell polarization drives the cytoskeletal morphology changes upon optogenetic activation of stress fiber formation, and it is consistent with the fact that RhoA signaling complexes are most abundantly active in the cell rear during retraction^23–25^.

Light-dependent increases in stress fiber levels were also of large effect size (**Figure 2b**). The basal stress fiber extent was similar between BcLOV4-mCherry control, opto-RhoA, and opto-RhoGEF11, suggesting little leakiness in RhoA signaling from diffusive contact in the dark-adapted state, even at the over-expression levels supported by HEK cells. Likewise, the morphological parameters were largely unchanged upon illumination of the effector-less control cells. Cytoskeletal changes were largely abrogated in cells pre-treated with pharmacological inhibitors of RhoA-GEFs (Rhosin) and/or (Rho-associated protein kinase) ROCK signaling (with Y-27632) (**Supplementary Figure 6**), thus confirming the RhoA signaling-dependence and that dark-adapted opto-RhoA is GDP-bound. The kinetics of dynamic membrane association (τ_on-RhoA_ = 1.27 sec, τ_on-GEF11_ = 1.13 sec) and undocking (τ_off-RhoA_ = 114.2 sec, τ_off-GEF11_ = 108.4 sec), measured in the presence of Rhosin and Y-27632 to suppress RhoA signaling, was similar to those of effector-less BcLOV4-mCherry (**Supplementary Figure 7**).

**Supplementary Figure 6: Pharmacological suppression of optogenetic RhoA pathway signaling.**

**Supplementary Figure 7: Membrane translocation kinetics in HEK cells.**

By all observed measures, GTPase-level stimulation by opto-RhoA was consistently more effective than GEF-level stimulation by opto-RhoGEF11. However, the differences were modest (small effect size and marginal significance). No differences were observed in basal stress fiber level or subcellular distribution patterns, although opto-RhoA expressed at slightly higher levels (**Supplementary Figure 8**). Together, these findings suggest that the net turnover of GDP/GTP-bound RhoA is not rate-limiting. It should be noted, though, that nuanced differences in signal integration may still exist since GTPase-level control integrates the inputs of multiple GEFs, whereas GEF-level control can be subtype specific (as shown by others for Rac1^26^). For simplicity and experimental throughput, our further characterization focuses on opto-RhoA since it was generally more efficacious (including its ability to separate adherens junctions) and fewer tools exist for directly controlling RhoA GTPase than for its GEFs (**Supplementary Videos 1 and 2**).

**Supplementary Figure 8: Expression level distribution in transfected HEK cells.**

### Spatial determinants of RhoA induction and signaling

In cells responding to whole-field stimulation, the magnitude of the response was largest along the polarization axis with the trailing edge contracting toward the leading edge, which suggests that the cell polarization drove the vectorial change in cell shape. We next explored the effects of subcellularly patterned stimulation on RhoA-driven cell contraction, specifically whether the orientation of blue light stimulation with respect to the cell polarization axis affects the vectorial change in cell morphology. (**Figure 3**). We defined the “stimulation angle” as the angle between the polarization axis and a line segment between the cell centroid and the stimulated ROI (30° bins, ~10 μm box and ~20% fraction of membranes) (**Figure 3a**). The magnitude of induced change clearly trended with the stimulation angle alignment with the polarization axis (**Figure 3b-d**). Stimulation at the leading and trailing edges led to largest constrictions, which was expected based on the organization of the underlying stress fiber network and endogenous subcellular distribution of RhoA signaling complexes^23–25^. These experimentally determined spatial relationships, between the input of optogenetic RhoA signaling induction and the downstream output of cytoskeletal contractility, will be useful for guiding experimental design and data interpretation across optical stimulation paradigms.

**FIGURE 3:**
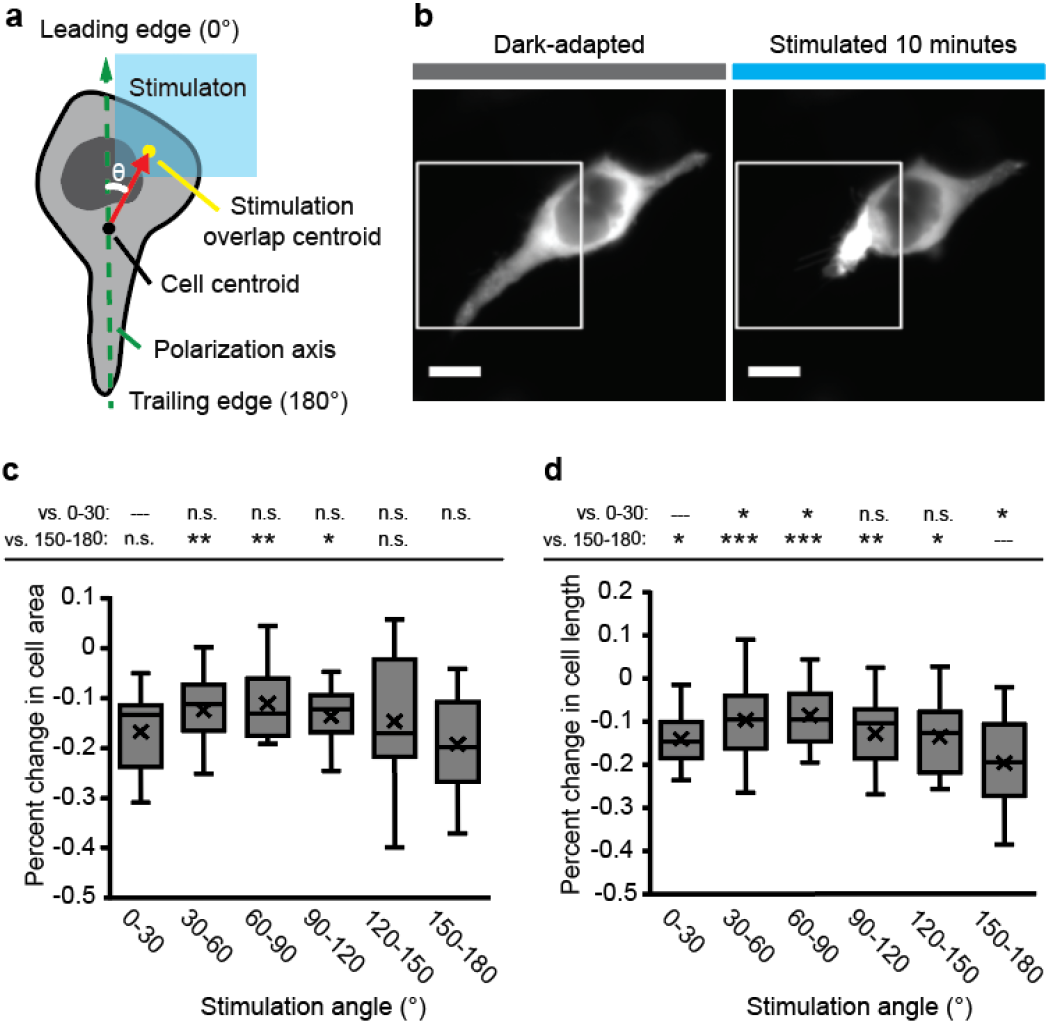
Stimulation angle dependence of opto-RhoA driven contraction. **a**. Schematic of stimulation angle (θ) calculation from the polarization axis, cell centroid, and centroid of the overlap region of the cell with the patterned stimulation field. **b**. Exemplar images of focal contraction of the trailing edge of a HEK cell after 10 minutes of pulsatile patterned stimulation (1.6% duty ratio). White box = illumination field. Scale = 10 μm. **c**. Box-and-whisker plot of change in cell area (relative to initial area) for binned stimulation angles. **d**. Box-and-whisker plot of change in cell length (relative to initial length) for binned stimulation angles. N = 10-35 independent videos per bin. Mann-Whitney U test: (*) p < 0.05; (**) p < 0.01; (***) p < 0.001; (n.s.) not significant. Top row = vs. 0-30° leading edge bin; bottom row = vs. 150-180° trailing edge bin.

### RhoA-driven mechanotransduction

One way that RhoA regulates cellular dynamics is mechanotransduction, the cytomechanical activation of gene expression (**Figure 4**). RhoA-controlled stress fibers relay mechanical cues from the extracellular maxtrix (ECM) to the transcriptional co-activator Yes-associated protein, YAP (and its paralog, TAZ) that regulates the activity of other transcription factors, like TEAD-family transcription factors^27,28^. Previous work by others showed that optogenetic activation of ARHGEF11 could drive rapid nuclear import of YAP within ~5 minutes of the increased cytoskeletal tension^9^. We report that Opto-RhoA activation drove nuclear import of GFP-tagged YAP (**Figure 4a-d**) on similarly fast timescales of ~3 minutes.

**FIGURE 4:**
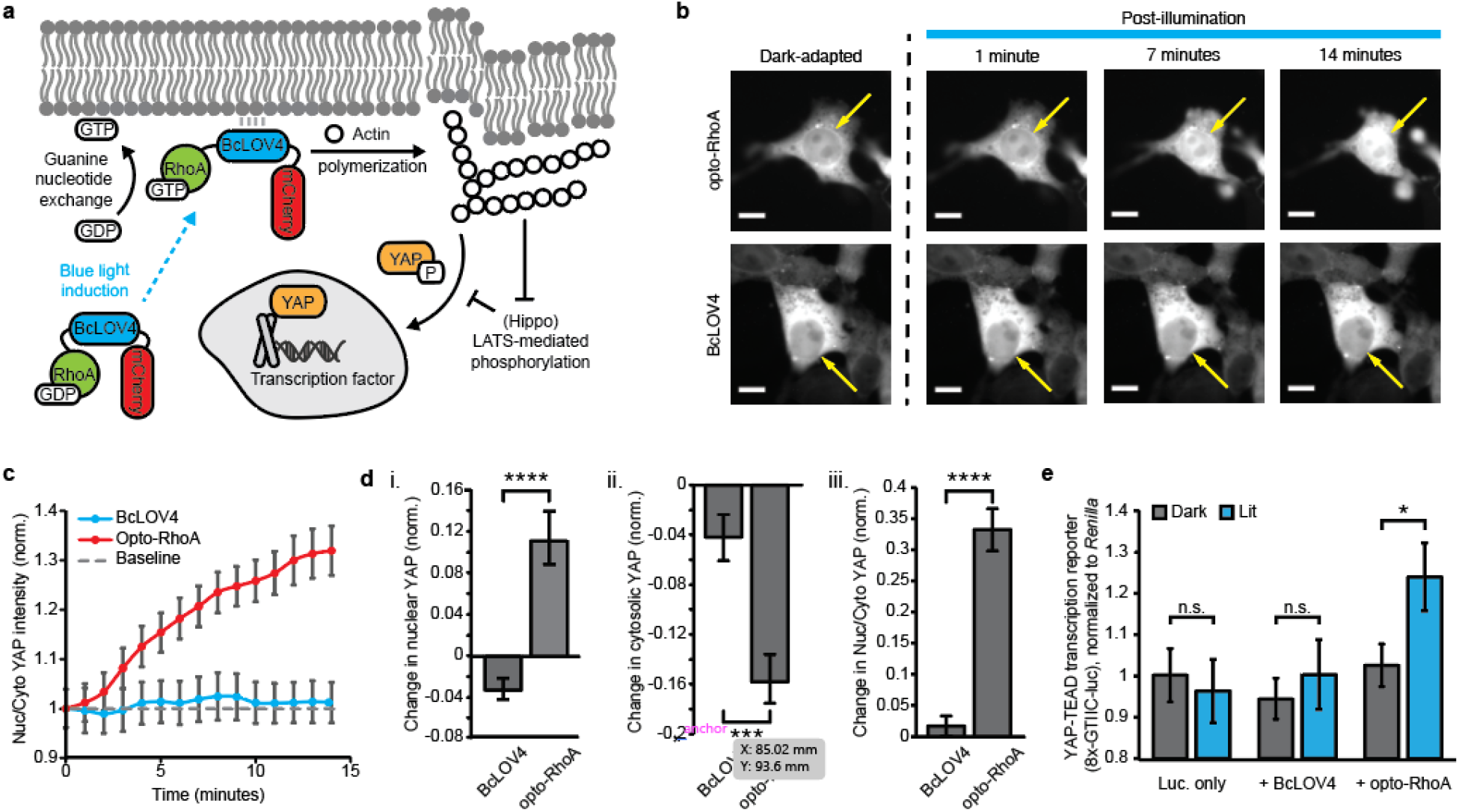
Opto-RhoA induction of YAP-dependent mechanotransduction. **a**. Photoactivated opto-RhoA drives actin polymerization and increases cytoskeletal tension to drive nuclear import of YAP and transcriptional co-activation. **b**. Exemplar images of YAP-GFP nuclear import following blue-light photoactivation of opto-RhoA and BcLOV4 control. Scale bar = 10 μm. Arrows indicate nucleus position. **c**. Nuclear:cytosolic YAP in response to 1.6% duty ratio stimulation of opto-RhoA and BcLOV4 control. N = 30 cells each. **d**. Change in (i) nuclear YAP, (ii) cytosolic YAP, and (iii) nuclear:cytosolic YAP, normalized to region area. N = 30 cells each. **e**. Dual luciferase reporter assay showing increased YAP-coactivated TEAD-dependent transcription driven by opto-RhoA but not BcLOV4-only control. Luminescence was measured from firefly luciferase under the regulation of a YAP-sensitive synthetic promoter (“8xGTIIC”), normalized to co-expressed *Renilla* luciferase. N = 8 wells per condition. **c-e**. mean +/- std err. Mann-Whitney U test: (*) p < 0.05; (***) p < 0.001; (****) p < 0.0001; n.s. not significant.

Beyond nuclear localization, we confirmed YAP/TAZ-TEAD-dependent transcriptional activation using a previously described^5^ luciferase reporter (**Figure 4e**) in serum-starved cells of low initial cytoskeletal tension. Thus, single-component optogenetic RhoA tools for driving downstream gene expression to study mechanotransduction that are complementary to recently reported optogenetic YAP/TAZ domains^29,30^ that shuttle into the nucleus when released from mitochondrial membranes into the cytosol by heterodimer dissociation. The RhoA-YAP relay is of particular importance in programming cell migration and tissue development^31–33^.

## DISCUSSION

We have expanded the repertoire of single-component optogenetic membrane recruitment-based tools to RhoA signaling, by fusing BcLOV4 to wildtype RhoA GTPase or the catalytic domain of ARHGEF11. We fused wildtype RhoA over constitutively active mutants to limit the probability of RhoA signal transduction in the dark by diffusive membrane contact alone, which was evident in cells expressing a BcLOV4-fused constitutively active G14V RhoA mutant with elevated basal stress fiber levels (**Supplementary Figure 9**). The dynamic range improvement by using wildtype GTPase was also observed with our previously reported opto-Rac1^22^.

Because the GTPase in opto-RhoA lacks a prenylation site^34,35^, it is less likely to be activated by the external stimuli that activate the endogenous cellular RhoA. Thus, the occlusion of the C-terminus of RhoA GTPase in our tool also enhances the orthogonality of opto-RhoA to the endogenous GTPase. In addition, opto-RhoA lacks a GDI binding site because of the disrupted prenlyation, thus preventing the over-expressed opto-RhoA from destabilizing the endogenous RhoA pool by otherwise outcompeting it for regulatory GDI interactions. These features are useful for leaving the basal cytoskeletal physiology intact when opto-RhoA is inactive in its dark-adapted state.

Opto-RhoA and opto-RhoGEF exhibit similar membrane association kinetics to BcLOV4, which we have previously demonstrated are favorable for manipulating cytoskeletal dynamics with high spatiotemporal precision. Throughout the data herein, opto-RhoA accumulation is observable in DMD-patterned illumination fields, even well after the induction period is complete. Such accumulation of activated Rho GTPase is also observed with a heterodimerizing optogenetic Cdc42 created by others^36,37^. However, it is not as readily observed in control cells inhibited with GEF and ROCK inhibitors, or in effector-less BcLOV4-mCherry controls, which exhibit similar membrane dissociation kinetics to one another. Together, these photoactivated distribution profiles and kinetics data suggest that the puncta originate *de novo* and persist from interactions amongst RhoA/actomyosin signaling complexes^38,39^, as opposed to LOV homo-oligomerization into large photobodies, which is the clustering mechanism of Cry2-RhoA we previously created.

In summary, we have created high-performance single-component tools for optogenetic activation of RhoA GTPase and RhoA-actvating GEFs to control cell contractility and RhoA-induced transcriptional mechanotransduction. The accompanying characterization of how cytoskeletal changes depend on spatial patterning of the optical stimulation informs how tool performance should vary across different experimental setups and designs, and will enable the study of diverse cell behaviors by connecting spatiotemporal RhoA signaling input to cytoskeletal output. These tools further demonstrate the versatility of BcLOV4 technology for single-component control over peripheral membrane proteins.

**Supplementary Figure 9: Basal and induced activity of constitutively active opto-RhoA-G17V (constitutively active) mutant.**

## MATERIALS AND METHODS ONLINE

### Genetic constructs

Domain arrangement combinations of BcLOV4, mCherry, and RhoA or GEF effector (with a flexible (GGGS)_2_ linker between each pair) were assembled by Gibson cloning using NEB HiFi DNA Assembly Master Mix (E2621) into the pcDNA3.1 mammalian expression vector under the CMV promoter. BcLOV4 and mCherry were amplified from their mammalian codon-optimized reported fusion (Addgene plasmid 114595)^20^. Wildtype RhoA GTPase was amplified from CLPIT Cry2PHR-mCherry-RhoA (Addgene plasmid 42959) without the C-terminal “CAAX” motif to prevent prenylation^18^. The DNA sequences of ARHGEF11 (Genbank ID XP_011508491.1) were human codon-optimized using the Integrated DNA Technologies (IDT) Codon Optimization Tool and ordered as gBlocks®. The DH domains of these GEFs, identified using the PROSITE ExPASy database, were amplified from these gBlocks. The RhoA constitutively active G14V mutant^40^ was generated by QuikChange site-directed mutagenesis. All genetic constructs were transformed into competent *E. coli* (New England Biolabs, C2984H). All sequences were verified by Sanger sequencing. For mechanotransduction assays, EGFP-tagged YAP was acquired from Addgene (plasmid 17843), and YAP-sensitive promoter plasmid was acquired from Addgene (plasmid 34615). Plasmids for opto-RhoA-mCherry and opto-RhoGEF-mCherry will be distributed through Addgene (plasmids 164472 and 164473).

### Optical hardware

Fluorescence microscopy was performed on an automated Leica DMI6000B fluorescence microscope under Leica MetaMorph control, with a sCMOS camera (pco.edge), an LED illuminator (Lumencor Spectra-X), and a 63× oil immersion objective. Excitation source illumination was filtered at the LED source (mCherry imaging λ = 575/25 nm; GFP or AlexaFluor488 imaging or wide-field BcLOV4 stimulation λ = 470/24 nm). mCherry was imaged with Chroma filters (T585lpxr dichroic, ET630/75 nm emission filter, 0.2-0.5 s exposure). Cells were imaged at room temperature in CO2-independent media (phenol-free HBSS supplemented with 1% l-glutamine, 1% penicillin-streptomycin, 2% essential amino acids, 1% nonessential amino acids, 2.5% HEPES pH 7.0, and 10% serum). The custom spatially patterned illuminator was (DMD) digital micromirror device-based and constructed from a digital light processor (DLP, Digital Light Innovations CEL5500), as previously described^22^.

### Mammalian culture and transduction

HEK293T (ATCC, CRL-3216) cells were cultured in D10 media composed of Dulbecco’s Modified Eagle Medium with Glutamax (Invitrogen, 10566016), supplemented with 10% heat-inactivated fetal bovine serum (FBS) and penicillinstreptomycin at 100 U mL^-1^. Cells were maintained in a 5% CO2 water-jacketed incubator (Thermo/Forma 3110) at 37 °C. Cells were seeded onto poly-d-lysine-treated glass bottom dishes (MatTek, P35GC-1.5-14-C) or into 24-well glass bottom plates (Cellvis, P24-1.5H-N) at 15-20% confluency. Cells were transfected at ~30-40% confluency 24 hours later using the TransIT-293 transfection reagent (Mirus Bio, MIR2700) according to manufacturer instructions. Cells were imaged 24-48 h post-transfection.

### Microscopy assays

#### Expression characteristics and membrane translocation

For membrane recruitment quantification, prenylated GFP was co-transfected as a membrane marker with the BcLOV4 fusions as previously described^20,22^. Briefly, an mCherry fluorescene image (500 ms exposure) was captured to assess protein expression level and subcellular distribution. Cells were then stimulated with a 5 s-long wide-field blue light pulse to stimulate BcLOV4, during which time mCherry fluorescence images were captured every 200 ms. The GFP marker was imaged immediately after blue light stimulation. For membrane dissociation via thermal reversion of the protein, cells were incubated with RhoGEF inhibitor Rhosin (Millipore-Sigma 555460) at 25 μM and ROCK inhibitor Y-27632 (Millipore-Sigma Y0503) at 10 μM for 24 hours prior to imaging to prevent cell contraction. mCherry was visualized every 5 s for 10-15 minutes in the absence of blue light stimulation. Membrane localization and dissociation were measured by line section analysis and correlation with prenylated GFP in ImageJ and MATLAB as previously described^20^.

#### DMD stimulation imaging

mCherry fluorescence was imaged every 15 s for 10 min. During this time, cells were stimulated with one second per minute (1.6% duty cycle) with patterned illumination of a 25 μm-wide square (~25% of cell area). The stimulation angle was defined as the angle defined by the polarization axis and the line segment between the cell centroid and the centroid of the stimulated cell area.

#### Pharmacological inhibitors

The GEF inhibitor Rhosin (Millipore-Sigma 555460) was added to cells at a final concentration of 25 uM following transfection, 24 hours before imaging. ROCK inhibitor Y-27632 (Millipore-Sigma Y0503) was added to cells at a final concentration of 10 uM at transfection, 24 hours before imaging.

#### Phalloidin staining

24 hours after transfection, cells were washed with PBS and media was replaced with DMEM supplemented with penicillin-streptomycin without FBS. Light-exposed plates were incubated under blue LEDs strobing at a 1.6% stimulation duty cycle in a 5% CO2 water-jacketed incubator for two hours. Cells were fixed with 4% paraformaldehyde in PBS at room temperature for 10 minutes, washed twice with PBS, then permeabilized with 0.1% Triton-x-100 in PBS for 15 minutes. Cells were blocked with 1% BSA in PBS for 30 minutes, then stained with Alexa Fluor 488 Phalloidin (Invitrogen, A12379) diluted 1:400 in PBS. Plates were washed twice prior to imaging. Total filamentous actin was quantified by normalizing total cell fluorescence at 488 nm to cell area, with N = 40 cells per condition.

#### RhoA-driven YAP mechanotransduction

For YAP nuclear translocation assays, HEK293T cells were initially plated at 75% confluency in 10 cm dishes to drive YAP to the cytosol. Cells were then passaged one day later and seeded at 25% confluency in 35 mm poly-d-lysine-treated glass bottom dishes. The next day, dishes were washed with PBS and media was replaced with DMEM supplemented with 2% heat-inactivated FBS and penicillin-streptomycin, then cells were transfected with a 4:1 ratio of opto-RhoA-mCherry or BcLOv4-mCherry to EGFP-YAP.

Over a 15 minute time-course, EGFP-YAP was imaged every 15 seconds using a 250 msec excitation light pulse that also stimulated BcLOV4, and mCherry fluorescence was imaged every minute. Nuclear and cytosolic fluorescence normalized to each region’s area was calculated at each minute by manual segmentation in ImageJ for N = 20 cells per condition.

For transcriptional mechanotransduction assays, we quantified YAP-TEAD dependent transcriptional activity using a dual luciferase reporter system (Promega, E1910). HEK293T cells were co-transfected with a YAP-sensitive promoter driving firefly luciferase expression (8XGTIIC-luciferase, Addgene #34615), *Renilla* luciferase, and BcLOV4-mCherry or opto-RhoA-mCherry. At transfection, full media was replaced with DMEM supplemented with 2% heat-inactivated FBS and penicillin-streptomycin. Half the cells were incubated under pulsing blue light with a 1.6% stimulation duty cycle for 12 hours. Cells were lysed according to manufacturer instructions. Luminescence was measured in white 96-well plates (Corning, 3917) on a Tecan M200 spectrophotometer with a 10 second integration time. The firefly luminescence value for each sample was normalized to its *Renilla* luciferase readout. N = 8 lysate samples per condition.

### Data analysis pipeline

Change in cell area, cell length, and centroid movement were computed via a custom analysis Python script, as schematized in Figure 3. For whole-field stimulation assays, videos of an entire field-of-view were cropped so that each contained only one cell. Contours of the cell membrane and nucleus were identified using the threshold function in OpenCV. Each video was rotated by increments of 5 degrees until the cell’s long polarization axis aligned with the y-axis, and then a cell-bounding rectangle was calculated using the Imutils package with the short edge aligned with the x-axis; the angle of rotation was chosen such that the cell nucleus was closer to the top of the rectangle as a morphological marker for the cell leading edge. Binary masks of the cell at initial (t = 0 min) and final (t = 10 min) timepoints were created for each cell. The change in cell area was calculated by finding the percent change in the area bounded by the cell’s contours at the final timepoint relative to its area at the initial timepoint. Change in cell length was calculated using the height of the bounding rectangle at the final and initial timepoints.

For experiments with spatially confined illumination, a similar imaging processing workflow was followed. A mask of the stimulation region was created using the same thresholding function as before, and this mask was also rotated at the same angle as the rest of the masks. To calculate the angle of stimulation, masks of the cell region within and outside the stimulation zone at the initial and final timepoints were created using the OpenCV bitwise operation “and” and “xor” functions, respectively. The centroids of the overlap mask and whole cell were computed at the respective timepoints. The angle of stimulation was calculated as the angle between (i) the vector connecting the initial cell centroid and overlap region centroid and (ii) the vertical vector between the initial cell centroid and the leading edge.

### Actin imaging

HEK293T cells were co-transfected with a nuclear marker (mTagBFP-Nucleus-7) and miRFP703-tagged LifeAct. The data analysis pipeline was applied to align the long axis of the cell with the y-axis, and to position the nucleus closer to the top of the cell. The actin arc was then located by identifying the region of brightest actin staining at least 500 pixels in area. The angle between the long axis and the vector connecting the cell centroid to the actin arc centroid was then calculated to verify that the actin arc occurs at the computationally identified leading edge of the cell.

### Statistical analysis

For whole-field stimulation assays, each cell was treated as a separate data point, with N = 82-93 cells from 10-12 field-of-view videos per condition. For spatially patterned stimulation assays, each data point was derived from one cell in an independent video. Cells were binned by angle of stimulation, with bin widths of 30 degrees spanning 0-180 degrees, N = 10-35 videos per bin. Statistical significance was assessed by the non-parametric Mann-Whitney U test, uncorrected for multiple comparisons.

## AUTHOR CONTRIBUTIONS

EEB designed genetic constructs, designed all experiments, and conducted all experiments. IAK created the patterned illumination system and assisted with the automated data analysis pipeline development. KY assisted with genetic construct design, engineering, and assays. BYC, JDB, and LJB coordinated all research. All authors contributed to experiment design, data analysis, and manuscript preparation.

## ACKNOWLEDGEMENTS

BYC acknowledges the support of National Science Foundation (NSF) Systems and Synthetic Biology (MCB 1652003), NIH/National Institute on Drug Abuse (R21 DA040434), and NIH/National Institute of Neurological Disorders and Stroke (NINDS) (R01 NS101106). EEB acknowledges the fellowship support of the National Institute of Neurological Disorders and Stroke of the National Institutes of Health (T32 NS091006). IAK acknowledges the fellowship support of the Paul and Daisy Soros Fellowship for New Americans and the NIH/National Institute of Mental Health F30 Award. LJB acknowledges the support of the NIH/National Institute of General Medical Sciences (R35 GM138211, R21 GM 132831). JDB acknowledges support from the National Science Foundation, Division of Civil, Mechanical, and Manufacturing Innovation (CMMI 1548571) and the NIH/National Institute of Arthritis and Musculoskeletal and Skin Diseases (R01 AR073809).

