## Supplementary Figures for "Single-component optogenetic tools for inducible RhoA GTPase signaling"

Supplementary Figure 1: Single-component optogenetic control over RhoA signaling by GEF activation.

Supplementary Figure 2: Molecular engineering of opto-RhoA.

Supplementary Figure 3: Molecular engineering of opto-RhoGEF.

Supplementary Figure 4: Optogenetic induction of contractility by a single pulse of unpatterned stimulation.

Supplementary Figure 5: Actin imaging of HEK cell polarization axis.

Supplementary Figure 6: Pharmacological suppression of optogenetic RhoA pathway signaling.

Supplementary Figure 7: Membrane translocation kinetics in HEK cells.

Supplementary Figure 8: Expression level distribution in transfected cells.

Supplementary Figure 9: Basal and induced activity of constitutively active opto-RhoA-G17V (constitutively active) mutant.

Supplementary Video 1 and Supplementary Video 2**:** Opto-RhoA drives adherens junction separation in HEK293T cells.

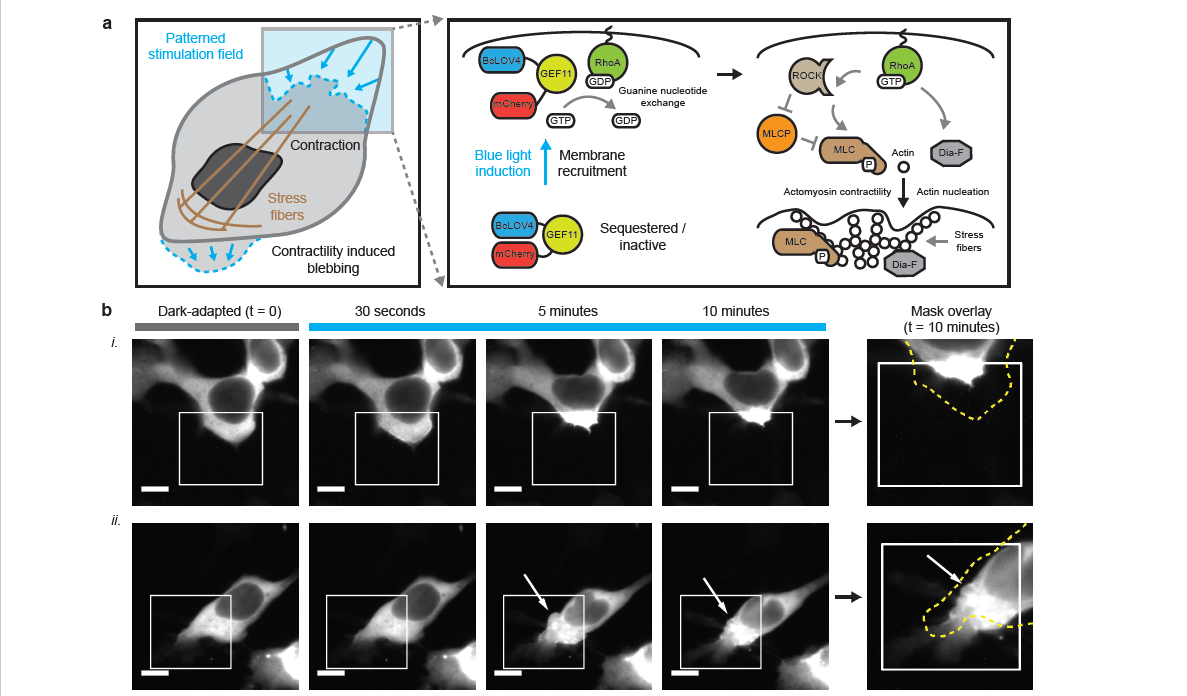

**SUPPLEMENTARY FIGURE 1**: Single-component optogenetic control over RhoA signaling by GEF activation. **a**. Schematized induction of cytoskeletal changes and contractile signaling in response to opto-RhoGEF11 activation by BcLOV4-mediated membrane translocation. **b**. Epifluorescence micrographs of HEK cells expressing opto-RhoA, visualized by mCherry. i. Leading edge contraction. ii. Contraction and reversible bleb formation. White arrow = bleb location. White box = spatially patterned blue light illumination field, stimulated at 1.6% duty ratio. Dotted yellow line = cell boundary mask in the dark-adapted state. Scale = 10 μm.

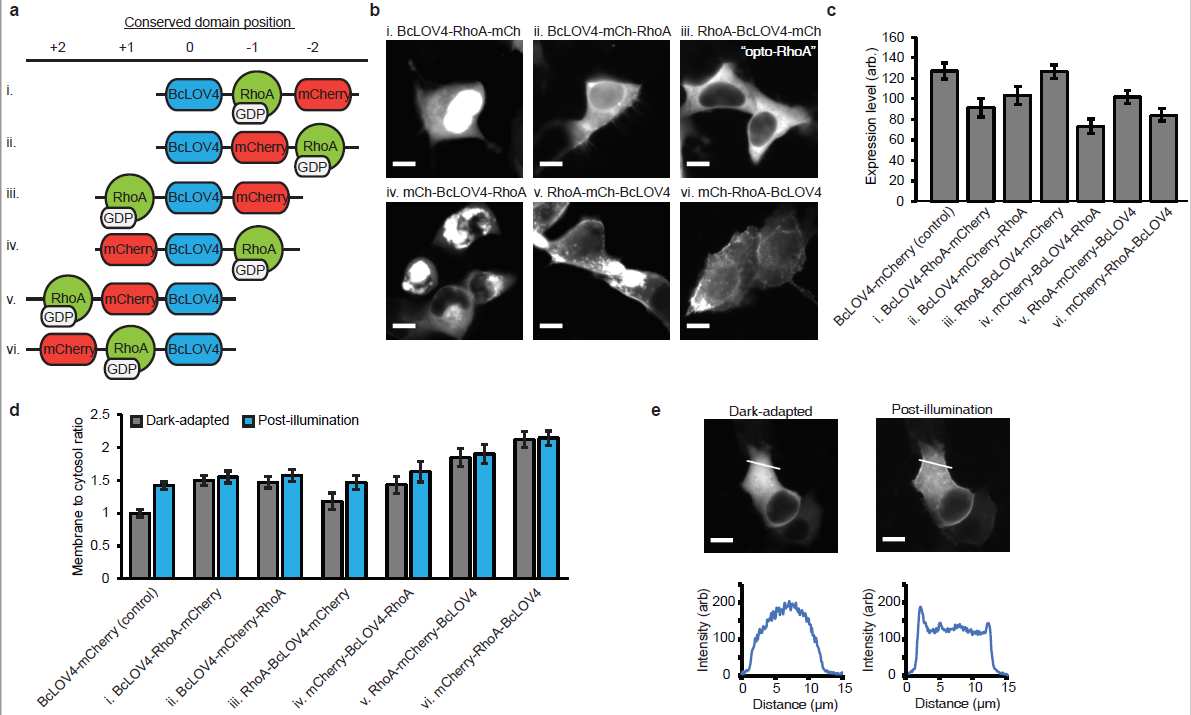

**SUPPLEMENTARY FIGURE 2**: Molecular engineering of opto-RhoA. **a**. Domain arrangement combinations of BcLOV4, wildtype (GDP-bound) RhoA GTPase, and mCherry visualization tag. Domains were separated by flexible (GGGS) 2 linkers. **b**. Fluorescence micrographs showing representative expression patterns of the six domain arrangements in the dark-adapted state in HEK293T cells. Scale = 10 μm. **c**. Relative expression level of genetic constructs versus BcLOV4-mCherry control with no effector. N = 25-31 cells per condition. Mean +/- std err. **d**. Ratio of membrane-localized vs. cytosolic protein for each domain arrangement in the dark-adapted and blue light-illuminated states, normalized to BcLOV4-mCherry control. N = 25-31 cells per condition. Mean +/- std err. **e**. Representative membrane localization of opto-RhoA following blue light stimulation. (Top) Fluorescence micrograph; scale = 10 μm. (Bottom) Line section pixel intensity.

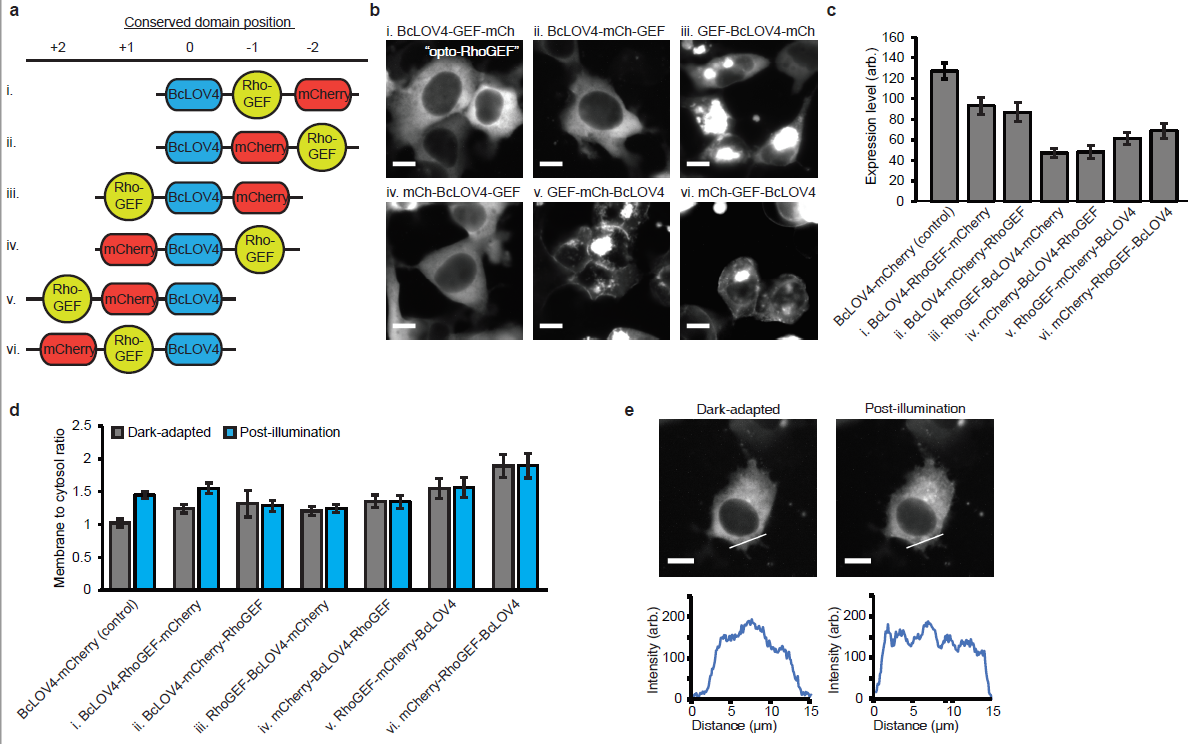

**SUPPLEMENTARY FIGURE 3**: Molecular engineering of opto-RhoGEF. **a**. Domain arrangement combinations of BcLOV4, the DH domain of ARHGEF11, and mCherry visualization tag. Domains were separated by flexible (GGGS)2 linkers. **b**. Fluorescence micrographs showing representative expression patterns of the six domain arrangements in the dark-adapted state in HEK293T cells. Scale = 10 μm. **c**. Relative expression level of genetic constructs versus BcLOV4-mCherry control with no effector. N = 25-31 cells per condition. Mean +/- std err. **d**. Ratio of membrane-localized vs. cytosolic protein for each domain arrangement in the dark-adapted and blue light-illuminated states, normalized to BcLOV4-mCherry control. N = 25-31 cells per condition. Mean +/- std err. **e**. Representative membrane localization of opto-RhoGEF following blue light stimulation. (Top) Fluorescence micrograph; scale = 10 μm. (Bottom) Line section pixel intensity.

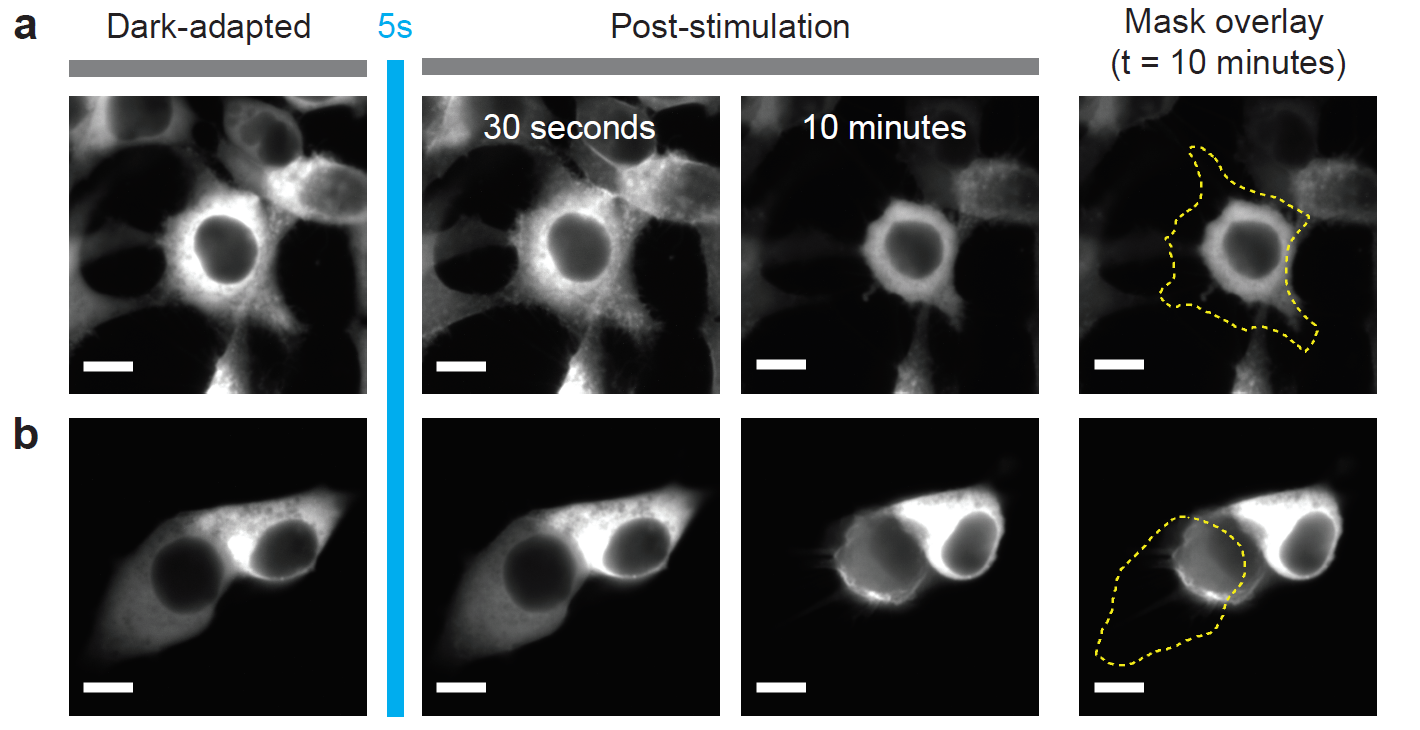

**SUPPLEMENTARY FIGURE 4**: Optogenetic induction of contractility by a single pulse of unpatterned stimulation. Representative images of HEK cells expressing (**a**) opto-RhoA and (**b**) opto-RhoGEF, before and after one 5-second pulse of wholefield blue light stimulation. Visualized by mCherry tag. “Mask Overlay” shows the initial cell boundary in yellow. Scale = 10 μm.

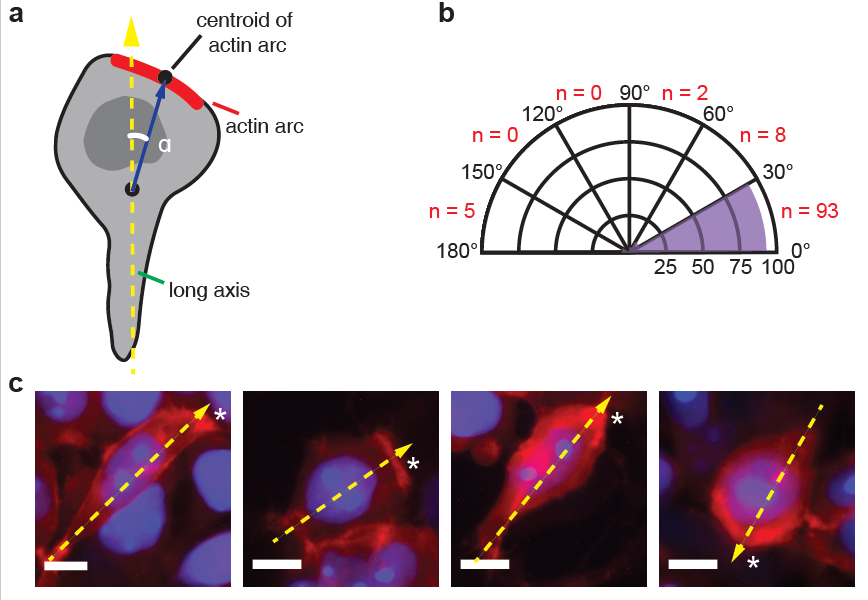

**SUPPLEMENTARY FIGURE 5**: Actin imaging of HEK cell polarization axis. **a**. The angle α is defined as the angle between the cell’s long axis (when the nucleus is positioned toward the top) and the line segment connecting the cell centroid to the centroid of the brightest region of F-actin, imaged by LifeAct-miRFP703. **b**. In wildtype HEK cells, the angle was overwhelmingly (in 93 of 108 cells, 86%) between 0 and 30 degrees, suggesting that the nucleus position along the long axis is a viable morphological marker to define the leading edge during automated analysis. **c**. Representative images of HEK293T cells expressing LifeAct (red) and a nuclear marker (mTagBFP-nucleus-7, blue). Yellow dotted line = algorithm-identified polarization axis of the cell. (*) = actin arc. Scale = 10 μm.

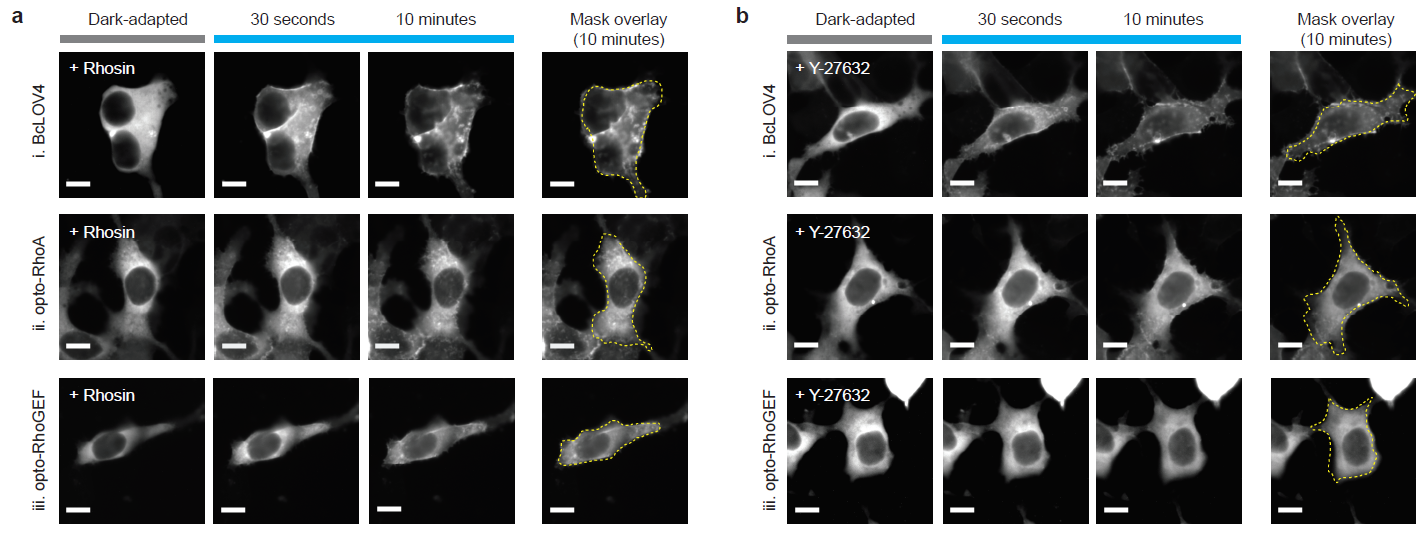

**SUPPLEMENTARY FIGURE 6**: Pharmacological suppression of optogenetic RhoA pathway signaling. **a**. Inhibition of RhoA-GEF interaction with Rhosin in representative cells expressiong (i) BcLOV4 control, (ii) opto-RhoA, (iii) opto-RhoGEF. Mask overlay shows the initial cell boundary (dotted yellow line). Optogenetic induction of contractility is largely abrogated. **b**. Inhibition of ROCK signaling by Y-27632 results in similar suppression. 1.6% stimulation duty ratio. Scale = 10 μm.

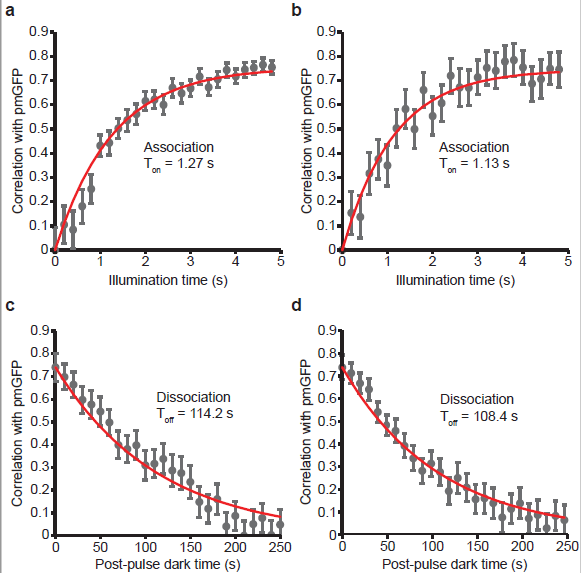

**SUPPLEMENTARY FIGURE 7**: Membrane translocation kinetics in HEK cells. Time constants were determined by correlation analysis between the membrane marker (pmGFP) and line

section profiles of the mCherry tag. RhoA signaling was pharmacologically inhibited with Rhosin and Y-27632 to isolate the membrane association/dissociation time constants attributable to the protein-lipid interaction. **a**. Membrane association of opto-RhoA-mCherry: T_on_ = 1.27 s, 95% CI 1.14-1.40 s. **b**. Membrane association of opto-RhoGEF-mCherry: T_on_ = 1.13 s; 95% CI, 1.01-1.26 s. **c**. Membrane dissociation of opto-RhoA-mCherry: T_off_ = 114.2 s; 95% CI, 106.6-121.8 s. **d**. Membrane dissociation of opto-RhoGEF-mCherry: T_off_ = 108.4 s; 95% CI, 103.4-113.4 s.

N = 20 cells per condition. Mean +/- std err. Values are in line with those of effector-less BcLOV4 in HEK and yeast cells, and of purified recombinant BcLOV4 with in vitro lipid interfaces.

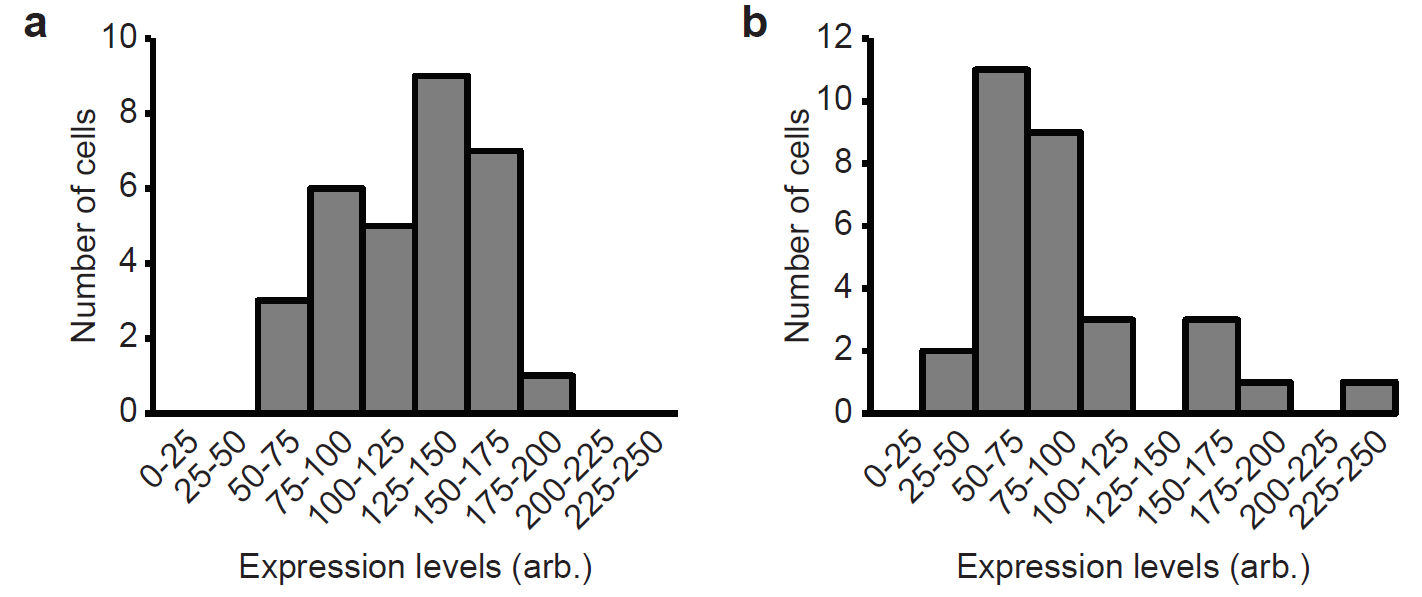

**SUPPLEMENTARY FIGURE 8**: Expression level distribution for transfected cells. (**a**) opto-RhoA and (**b**) opto-RhoGEF. N = 30-31 cells per construct.

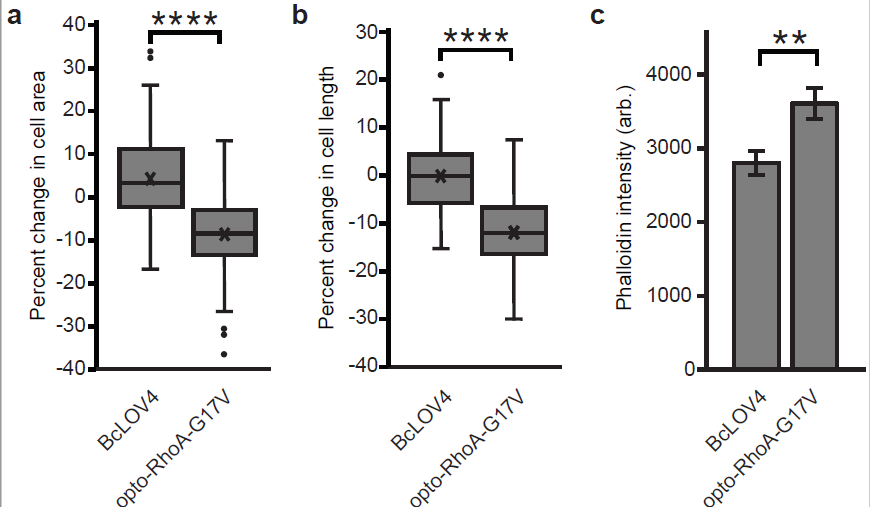

**SUPPLEMENTARY FIGURE 9**: Basal and induced activity of constitutively active opto-RhoA-G17V (constitutively active) mutant. Box-and-whisker plots of (**a**) change in cell area normalized to initial area, and (**b**) change in cell length, normalized to initial length, vs. BcLOV4-mCherry control (reproduced from main text Figure 3). N = 82-90 cells each. **c.** Quantification of basal actin stress fibers in transfected cells in the dark. Mean +/- standard error. N = 40 cells per condition. **a**-**c**. Mann-Whitney U test: (**) p < 0.01; (****); p < 0.0001; (n.s.) not significant.

**SUPPLEMENTARY VIDEO 1 and SUPPLEMENTARY VIDEO 2:** Opto-RhoA drives adherens junction separation in HEK293T cells. mCherry imaged every 15 seconds. White box = Patterned blue-light stimulation region (1.6% duty ratio). Scale = 10 μm.
